# Double-Stranded RNA Profiling with Mass Photometry

**DOI:** 10.64898/2026.05.15.725554

**Authors:** Matthew J. Ranaghan, Nathanial E. Clark, Kayleigh Fay, Alexander R. O’Shea, Natalia Markova, Svea Cheeseman, Gabriella Kiss

## Abstract

Double-stranded RNA (dsRNA) is a potent immunogenic impurity and its detection is a critical quality attribute in characterizing mRNA therapeutics. Standard analytical methods are only able to resolve the bulk presence of dsRNA and cannot characterize the different sub-species that may be present within a mRNA sample. In this study, we use mass photometry (MP) as a single-molecule analytical platform for the simultaneous detection and characterization of dsRNA impurities in mRNA samples. We demonstrate how ionic strength can interfere with the stability of the mAb/dsRNA complex and measure the binding affinity (1 nM) under a set of parameters for reproducible characterization of the complex. We leverage the J2 antibody to identify antibody/dsRNA complexes that resolve dsRNA-positive species within an mRNA sample based on discrete molecular weights. Furthermore, we introduce a novel MP assay that harnesses the repulsive surface chemistry of uncoated glass to exclude the bulk mRNA analyte for higher loading concentrations to sensitively profile trace dsRNA impurities as antibody-bound species. The semi-quantitative MP aligns with the bulk values determined for ELISA and dot blot detection of dsRNA. This work establishes MP as a valuable next generation mRNA analytical tool for analyzing dsRNA byproducts within mRNA samples.

**GRAPHICAL ABSTRACT:** 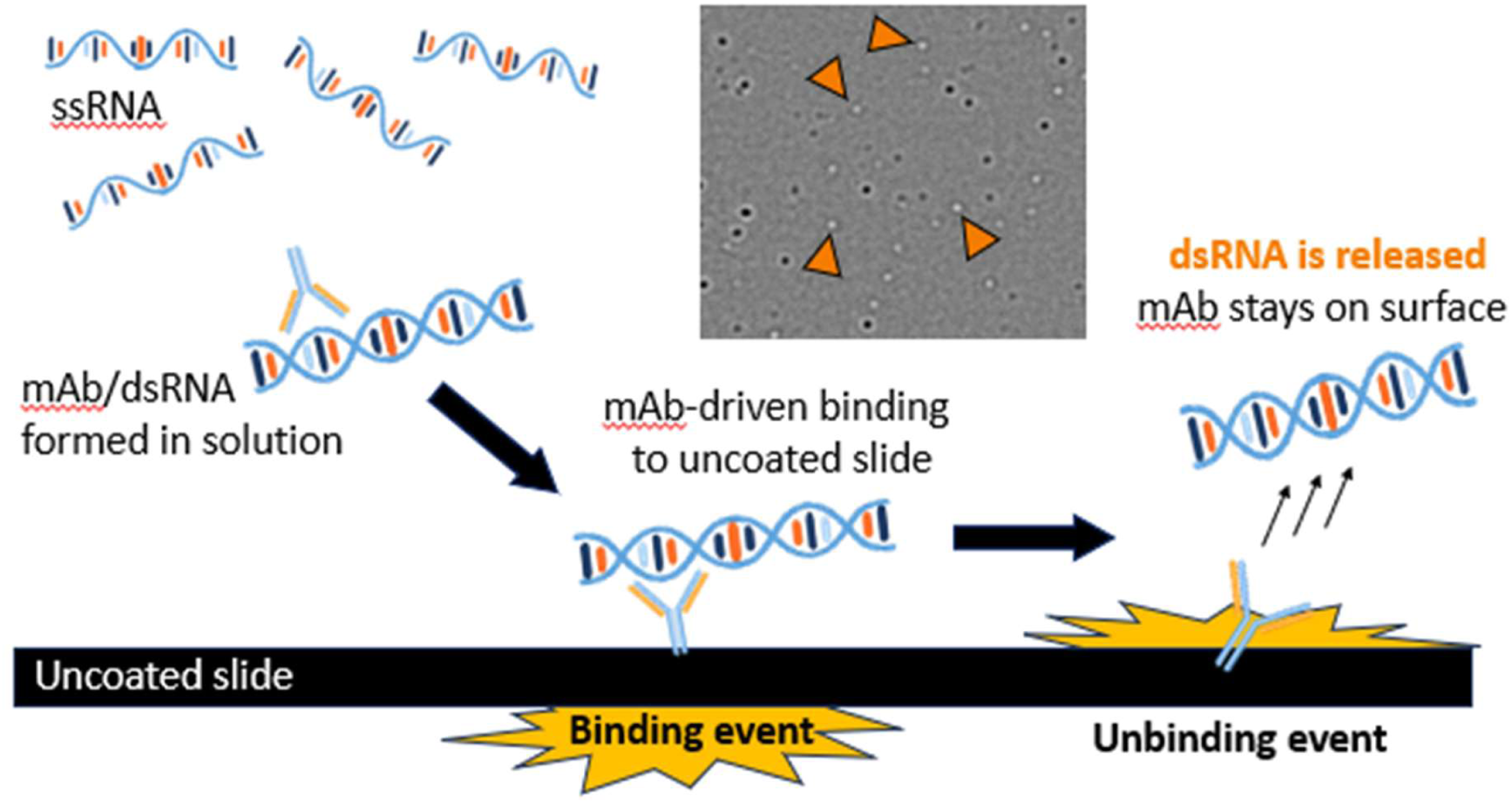

## INTRODUCTION

The clinical translation of *in vitro* transcribed (IVT) messenger RNA (mRNA) has fundamentally altered the landscape of genomic medicine, yet the maturation of the platform hinges on resolving a critical manufacturing bottleneck: the co-production of double-stranded RNA (dsRNA) impurities. While IVT enables the scalable synthesis of mRNA, it simultaneously generates process-related byproducts that are chemically identical to the therapeutic payload but adopt conformations that could be highly immunogenic, double-stranded conformations (1). Long dsRNA serves as a potent trigger for pattern recognition receptors (e..g, TLR3, RIG-I, and MDA5) (2–4). In a therapeutic context, this engagement drives robust type I interferon and pro-inflammatory cytokine responses; while potentially adjuvant-like for vaccines, unintended signaling can suppress mRNA translation and severely limit therapeutic protein yield (5, 6).

Achieving high purity mRNA (≥99.9%) remains a formidable engineering challenge because dsRNA and single-stranded mRNA are virtually indistinguishable by standard chemical means. These impurities arise through diverse mechanisms, including aberrant T7 polymerase activity, self-annealing, or template-derived duplex formation (7). While recent advances in dsRNA-affinity matrices and ion-pair reverse-phase HPLC (IP-RP-HPLC) have improved removal strategies (8, 9), the field lacks the analytical precision to characterize low-abundance, heterogenous dsRNA species. Existing benchmarks, such as the J2 mAb ELISA (10), are frequently confounded by matrix effects and false-positive signals from native mRNA hairpins. Consequently, batch-to-batch variability and systemic reactogenicity remain persistent hurdles for regulatory approval (1, 11). Hence there is an urgent need for orthogonal and complimentary high-resolution analytical modalities capable of defining the dsRNA profile to ensure the safety and potency of next-generation mRNA therapeutics.

Mass photometry (MP) is a single-molecule technique that is becoming popular for characterizing proteins (12), macromolecular complexes (13), nucleic acids (14, 15), and AAVs (16). Building on these successes, researchers are now using MP in mRNA workflows to determine sample integrity, aggregation, and batch-to-batch reproducibility (7, 17–21). *De Vos* and colleagues have demonstrated the feasibility of using MP to detect J2-mAb bound dsRNA on cation coated slides (22); however, they only present proof-of-concept data around purified standards. We present here a comprehensive MP-based assay to detect and characterize the various dsRNA impurities in mRNA samples, establishing MP as an essential tool for developing mRNA therapeutics.

## MATERIAL AND METHODS

### Reagents and chemicals

The J2 antibody (GenScript) was used to detect dsRNA. A 400 bp dsRNA standard (ACROBiosystems) was used as a defined dsRNA reference material. GFP mRNA (Aldevron) was purchased and processed for dsRNA impurities in accordance with Clark *et al*. (8). Aliquots for the input, flow through (FT), and elution were sampled and frozen until use. All RNA samples were stored in nuclease free water (New England Biolabs). All buffer preparations were carried out using 1× phosphate-buffered saline (PBS; Gibco, 14-190-250) or Tris-EDTA (TE) buffer (pH 8) (Integrated DNA Technologies). Sodium chloride stock solutions (5M NaCl; Teknova) was used for ionic strength adjustment where required. All reagents were used as received unless otherwise specified.

### UV-vis spectroscopy

A Nanodrop Lite Plus (ThermoFisher) was used to quantify all RNA samples immediately after preparation. An s_260_ of 5,200,000 M^-1^ cm^-1^ was calculated for the 400 bp dsRNA standard. An s_260_ of 8,100,000 M^-1^ cm^-1^ was calculated for the GFP mRNA sample.

### ELISA and immuno-norther blots

The EasyANA dsRNA kit was used for dsRNA quantification (Vazyme). The immuno-northern blot and dot-blots were performed as previously described (23).

### Mass photometry measurements

All MP experiments were collected using a TwoMP (Refeyn Ltd) using either MassGlass Uncoated (MGUC) and MassGlass Nucleic Acid (MGNA) microscope coverslips (Refeyn Ltd). Length calibration of nucleic acids was supported by a dsRNA ladder (New England Biolabs) and single-stranded RNA (ssRNA) ladders (New England Biolabs). Mass calibration for was performed using the MFP1 mass standard (Refeyn Ltd.). Experiments were done in Normal mode with a Large Field of View. Movies were collected for 60 seconds. Data were fit using DiscoverMP (Refeyn, Ltd).

### Binding affinity (K_D_) measurements

The K_D_ of the J2 mAb for dsRNA was determined in TE buffer (pH 8) supplemented with 50 mM NaCl. Titration samples were prepared with 2 nM of the 400 bp dsRNA standard and 0.8 to 100 nM J2 mAb before incubating mixtures for 30 min at ambient temperature. Mass photometry data of each sample was then collected on MGNA slides in triplicate. Experiments were mass calibrated against MFP1. Data were then fit in DiscoverMP (Refeyn Ltd) by manual fitting of Gaussians to observed peaks. Event counts for the J2 mAb, dsRNA, or mAb/dsRNA complexes were extracted using 20 kDa windows centred on the mass of each species. Data were then fit using event counts for only the free dsRNA substrate or as the sequential addition of J2 mAbs to the dsRNA substrate. A quadratic equation was used to model the data fitting only the free dsRNA substrate

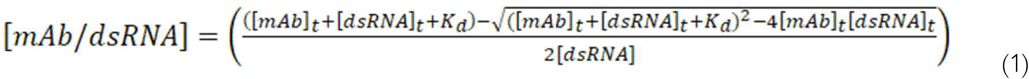

where the [mAb/dsRNA] represents the fraction bound population, [mAb]_t_ represents the total concentration of J2 mAb, [dsRNA]_t_ represents the concentration of dsRNA, and K_d_ is the apparent dissociation constant (24). For the kinetic model, the binding reaction forming a complex of mAb/dsRNA was modelled as the sequential addition of J2 mAb monomers to dsRNA:

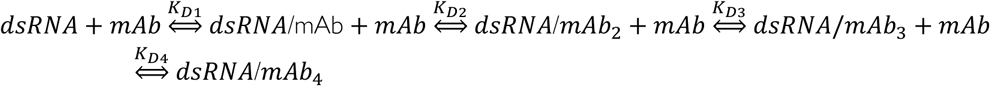

Generalising:

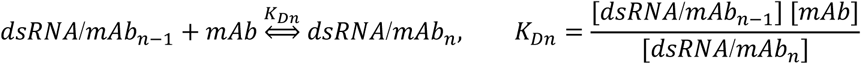

So:

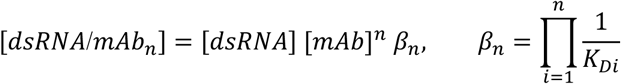

The total concentrations of dsRNA and mAb used at each point in the titration series are known ([*dsRNA*]_0_ and [*mAb*]_0_ respectively). Therefore, according to mass-balance:

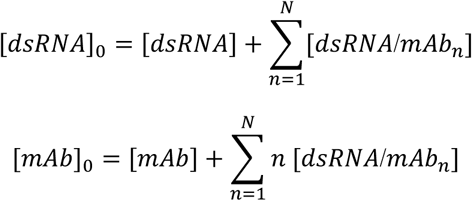

Thus:

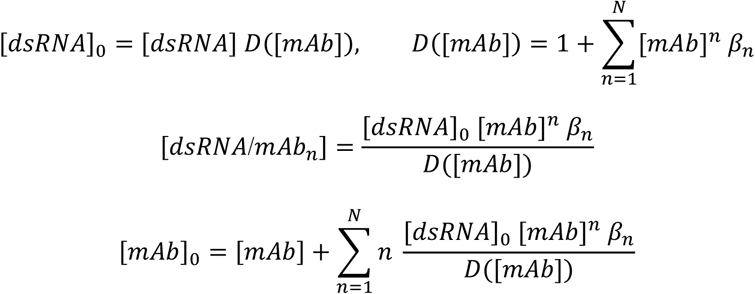

Let:

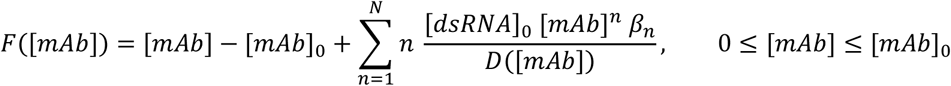

For each J2 mAb concentration used in the titration series, *F*([*mAb*]) = 0 was solved in its domain using Brent’s method and dsRNA/*mAb* _n_ complex concentrations were predicted (25, 26). Kinetic equilibrium constants were obtained by performing least-squares optimization between experimental species concentrations and those predicted by the model using the Trust Region Reflective algorithm (26, 27). Standard errors were calculated from the resulting Jacobian matrix.

### NaCl titration

Reagents were prepared at 2x in TE buffer with no supplemented NaCl. A serial dilution of NaCl concentrations was prepared at 1:1 in water at a 10x. Samples were prepared by adding equivalent amounts of J2 mAb and dsRNA to a tube preloaded with a 10x NaCl aliquot. Samples were then incubated for 30 min at room temperature before collecting MP data on a TwoMP. Samples replicates were collected for three independent experiments.

### MassFluidix detection of the mAb/dsRNA complex

Rapid dilution measurements were performed on a TwoMP with the MassFluidix HC system using uncoated MassFluidix HC Chips. MassFluidix data were mass-calibrated against manually diluted MP measurements of MFP1 of uncoated slides. The J2 mAb (2.5 uM) and mAb/dsRNA (1.25 uM for GFP mRNA or 150 nM for the dsRNA standard) complexes were prepared in TE buffer with 50mM NaCl and incubated at room temperature for 30 minutes prior to loading onto the MassFluidix HC system. Measurements were collected using the MassFluidix module in AcquireMP (Refeyn, Ltd) and Oxygen (Fluigent). Buffer flow rate was set to a constant 1 mL/min. Initial sample flow was set to 7.9 uL/min for a period of 90s, at which point flow was stopped, the sample tube was replaced with buffer, and flow was resumed. Upon reaching the observation window, the sample dilution rate was set to 1000x and the measurement was recorded for 60s. Data were fit using DiscoverMP (Refeyn, Ltd).

### Estimation of %dsRNA in mRNA fractions

Separation of dsRNA from GFP mRNA samples was performed in accordance with Clark *et al*. (2025) (8). Aliquots for the input, flow through (FT), and elution fractions were sampled and frozen until use. MP experiments were performed either on MGNA to detect complete mRNA profile or MGUC to detect the relative amount of dsRNA in the fraction. For MGNA detection, mRNA was prepared at 5 nM in IDTE buffer supplemented with 50 mM NaCl before collecting data in triplicate as described above. For MGUC detection of dsRNA species, mRNA samples were prepared at >100 nM in IDTE buffer supplemented with 50 mM NaCl before adding the J2 mAb (25 nM). Samples were incubated at ambient temperature for 30 min. MP data were then collected in triplicate for each sample as described above. The relative amount of dsRNA was then calculated by first fitting all MP data in DiscoverMP (Refeyn, Ltd.). Data collected on MGNA were fit by Gaussian distribution for the intact monomer and then interval gates were assigned to quantify counts for degradation (<200 kDa) and aggregate species (>300 kDa). Data collected on MGUC were first fit by Gaussian distributions to assign masses for identifiable populations in the binding and unbinding mass distributions. Gating was then assigned to quantify unbinding counts for degradation and aggregate species to avoid binding counts from the free J2 mAb at 150 kDa. Calculation of the %dsRNA content by MP was done by first normalizing all counts to 100 nM total RNA and mass-weighting counts. The relative %dsRNA was then determined as the ratio of unbinding counts from uncoated slides over the total counts from binding counts on MG-NA (coated) slides. For dot-blot quantification, the intensities of each spot were determined using ImageLab software (version 6.1.0, Bio-Rad Laboratories), and plotted against the mass of RNA contained in each spot to generate standard curves. Separate standard curves were prepared for the 50 mM NaCl and 150 mM NaCl samples. Statistical significance of the immunoblot was determined using a two tailed T-test.

## RESULTS

### Mass photometry of dsRNA

Analytical methods capable of resolving the intact mass of dsRNA-containing species are presently lacking in the field, contrasting sharply with the high-fidelity standards established for single-stranded mRNAs. Recent publications have demonstrated that MP can resolve mRNA length within 5% of the calculated length using standard cationic surfaces (17, 18, 20); however, similar demonstrations for dsRNA have not yet been clearly demonstrated. We therefore used standard RNA buffers (i.e., PBS and TE) to compare GFP mRNA and a 400 bp dsRNA standard. Figure 1 illustrates how the measured lengths of intact mRNA monomer and dsRNA standard are all within 3% of the calculated length, but also demonstrate how buffer choice is a critical factor data quality for dsRNA. Specifically, the MP histogram of dsRNA in TE buffer shows a monodisperse population with minimal low molecular weight species and negligible unbinding compared to PBS buffer. Unbinding events, characteristically represented as “negative mass” in the MP histogram (Figure 1C,D), indicate poor surface interactions from the biomolecule and can significantly increase the baseline noise of the recording (28) and likely stems from the physiochemical differences of ssRNA and dsRNA that enable their separation by pH or salt gradients (8, 29). These results, paired with the broader standard deviation (σ) of the observed peak at ∼400 bp, indicate that PBS is not ideal for dsRNA measurements by MP.

**Figure 1.**
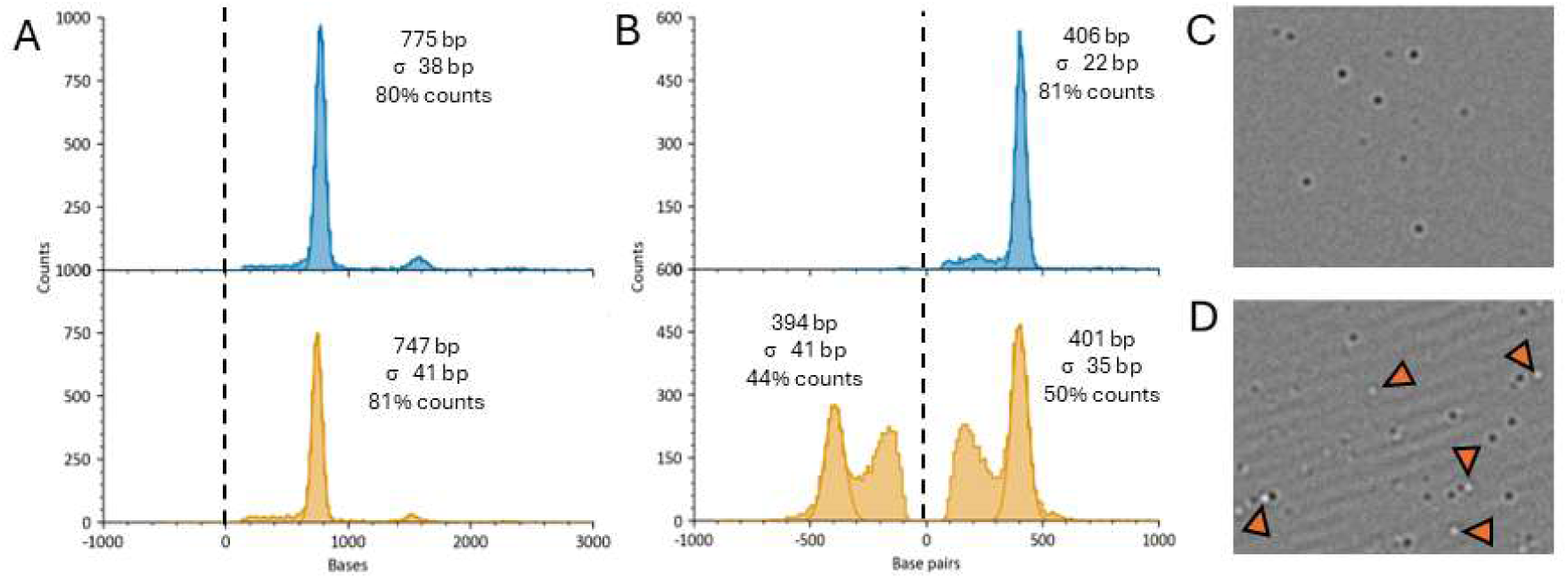
Effect of buffer on the MP profile of the RNA samples on MG-NA. (A,) Mass photometry histograms of GFP mRNA (750 b) in TE buffer (top) or PBS (bottom). The dotted line at the origin is a visual aid for separating the positive binding and negative unbinding masses. (B) Mass photometry histograms of the 400 bp dsRNA standard in TE buffer (top) or PBS (bottom). The dotted line is presented as described in A. (C) Field of view image for the 400 bp dsRNA standard in TE buffer. (D) Field of view image for the 400 bp dsRNA standard in PBS. Examples of unbinding events are denoted by orange triangles

The relative purity of RNA samples is also directly assessed in the MP data as either the relative counts from a given species or mass conversion to a weight-based value. The intact monomer of the mRNA samples is 80-81% counts in both TE and PBS (Figure 1A). This corresponds to an mRNA purity of ∼78% by weight and agrees well with the 76% purity reported by capillary electrophoresis in the manufacturer’s CoA document. This agreement also supports that mRNA characterization by MP is consistent across commonly used buffers and orthogonal techniques. Conversely, assessment of the dsRNA standard revealed a strong buffer dependence, whereby purity measurements in TE (81% counts or 87% weight) aligned more closely with the HPLC-derived CoA value (>90%) than by MP in PBS (50% counts or ∼50% weight) (Figure 1B). The high fidelity of the TE-based MP measurements relative to the manufacturer’s specifications identifies TE as the superior buffer for accurate MP assessment of dsRNA.

### Characterization of the J2 mAb/dsRNA complex

The J2 mAb is widely regarded as the gold-standard for detection of dsRNA containing species in biological samples. It selectively recognizes dsRNA segments in a sequence-independent manner, binding robustly to A form helices (14-40 bp) without deformation of the RNA and little reactivity to ssRNA or RNA/DNA hybrids (10, 30). This selectivity makes the J2 mAb a desirable reagent for applications that specifically detect dsRNA segments to include ELISA assays (10), immunofluorescence (31), and dot blotting (8), amongst others (32, 33).

We next sought to understand the interaction between the J2 mAb and dsRNA standard in TE buffer. Figure 2A shows how the J2 mAb and dsRNA substrate exist as well separated, monodisperse populations with no defined aggregate population in TE buffer. Mixture of the two reagents results in the generation of new peaks at 388 kDa and 550 kDa, which respectively represent a 1:1 and 2:1 complex of the mAb/dsRNA. The observed masses for all species are compared with calculated values in Table 1. Mass photometry typically reports measured length of RNA with <4% error. The larger relative error for the mass of dsRNA substrate is attributed to using an average nucleotide mass (660 Da per base pair) for an unknown sequence paired with differences in the refractive index of the calibrant used for these experiments (Supplementary Figure S1 and Supplementary Table S1). Specifically, a protein calibrant (MFP1) was chosen for all mAb-binding experiments because commercial ladders report nucleotide length of an unknown sequence and not absolute mass for the different species. Hence, we find it more accurate to calibrate against known protein standards for a relative mass of the dsRNA species and measurable iteration(s) of a bound antibody (i.e., addition of 150 kDa). For the mAb/dsRNA complex in Figure 2A, we indeed see a relative mass for the free dsRNA species (237 kDa) and two new populations for a dsRNA population with one bound (388 kDa) or two bound (544 kDa) antibodies. The relative error for measured masses of these species is noted to decrease with each additional mAb bound to the dsRNA.

**Figure 2.**
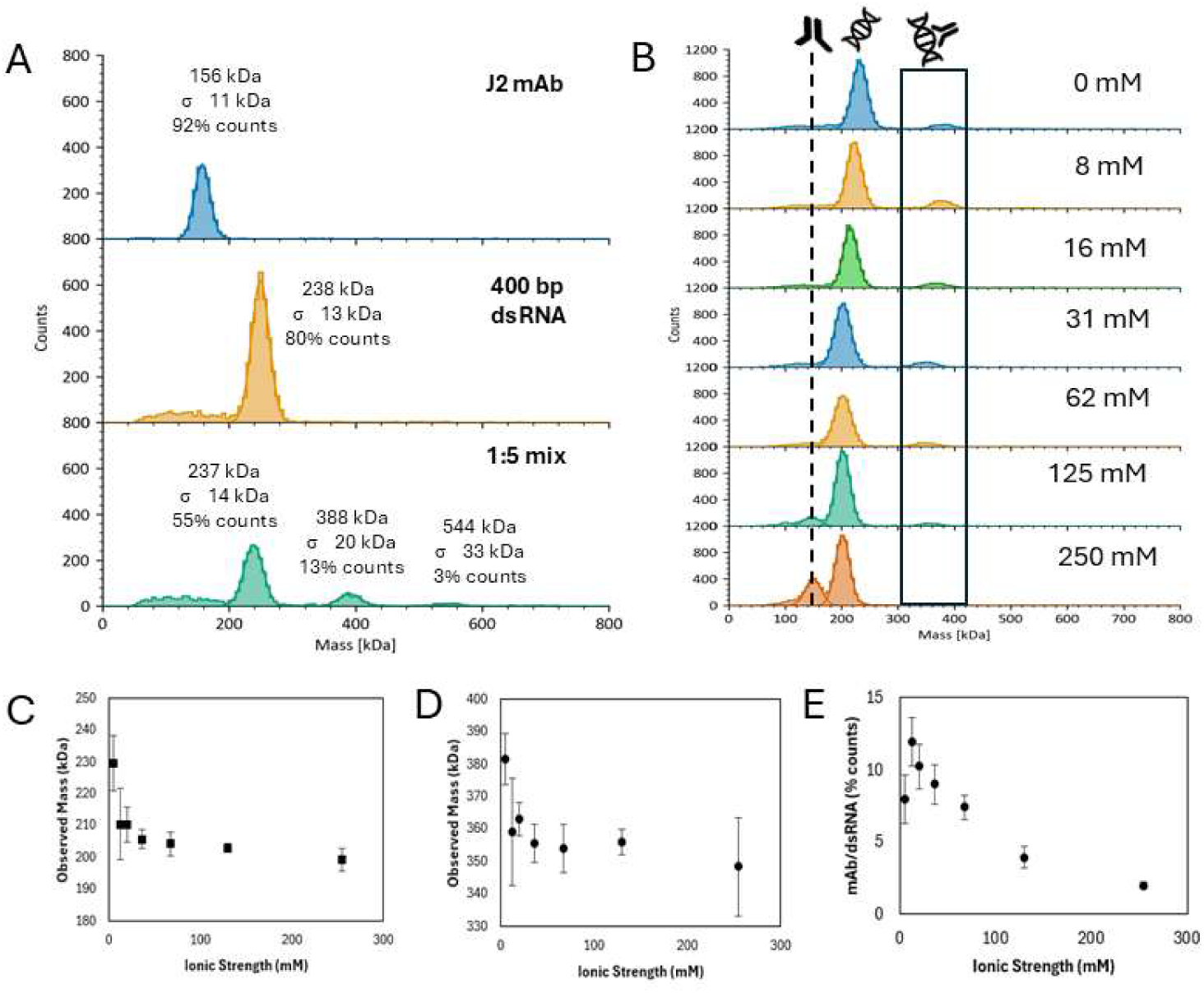
Characterization of the J2 mAb recognition of dsRNA. (A) Mass photometry histograms of the J2 mAb (*top*), 400 bp dsRNA (middle), and mAb/dsRNA complex (bottom). Samples were prepared in TE buffer individually or at a 1:5 ratio for the mAb/dsRNA complex. (B) Histograms of mAb/dsRNA samples prepared at a 1:5 ratio. Distinct peaks are observed for the free J2 mAb (150 kDa), free dsRNA (220-240 kDa), and mAb/dsRNA complex (340-390 kDa). All samples were prepared in TE buffer with supplemented NaCl to control ionic strength. (C) Average mass distribution of the free dsRNA with increasing ionic strength. (D) Average mass distribution of the mAb/dsRNA complex with increasing ionic strength. (E) Relative stability of the mAb/dsRNA complex as a function of ionic strength. Data represent the relative population of mAb/dsRNA complexes (350 kDa and larger) after normalization to experimental counts. Samples were prepared at a 1:5 ratio for mAb/dsRNA in TE buffer supplemented with NaCl. Error bars represent the standard deviation from experimental replicates.

**Table 1.** Calculated and observed masses for dsRNA-J2 mAb assemblies. Single-molecule mass photometry analysis of molecular weight deviations. Relative error percentages provide a benchmark of measurement accuracy compared to sequence-derived values. A clear reduction in mass error is observed for higher-order nucleoprotein complexes.

|  | Calc. mass (kDa) | Observed mass (kDa) | Relative error (% calc) |
| --- | --- | --- | --- |
| J2 mAb | 151 | 156 +/- 2 | 3.3 |
| 400 bp dsRNA | 272 | 249 +/- 2 | 8.5 |
| dsRNA + 1 mAb | 423 | 388 +/- 3 | 8.3 |
| dsRNA + 2 mAb | 574 | 544 +/- 9 | 5.2 |

The observed buffer effects led us to investigate the salt sensitivity of the mAb/dsRNA complex. To this end, we mixed the J2 mAb and dsRNA substrate at a 1:5 ratio to bias the complex to a single mAb per complex. Samples were assembled in TE buffer supplemented with an increasing amount of NaCl and allowed to reach equilibrium prior to MP measurements. The relative amount of mAb/dsRNA complex was then quantified and normalized against all binding counts (Figure 2B). A negative trend is observed for the measured mass of the dsRNA species and the measured mAb/dsRNA complex is observed at low ionic strength and stabilizes at ∼ 50 mM (Figure 2C,D). A similar negative trend is also observed for the relative abundance of the mAb/dsRNA complex with nearly complete inhibition at >100 mM ionic strength (Figure 2E). Such salt effects raise concern about quantification of dsRNA impurities across commercially available kits that use physiological buffer conditions (e.g., phosphate or tris buffered saline solutions).

Regardless, the net difference between the free dsRNA and mAb/dsRNA complex remains at 151 +/- 2 kDa at all ionic strengths. These data led us to supplement 50 mM NaCl into our TE buffer for all subsequent experiments to stabilize variability from “carry over” reagents in mRNA samples (*vide infra*).

The *K*_D_ for the J2 mAb interaction with the dsRNA substate was measured in TE buffer with 50 mM NaCl. Titration of the J2 mAb into a fixed amount of the dsRNA (2 nM) revealed an increasing number of species as the mAb concentration increased (Figure 3A). Each new species exhibited a mass of +150 kDa, which represents the sequential addition of a J2 mAb to the dsRNA substrate. The *K*_D_ for for the mAb/dsRNA complex was determined to be 3.6 +/- 0.4 nM by monitoring either the depletion of the free dsRNA (220 kDa) with increasing mAb concentration (Figure 3B) or 0.90 +/- 0.66 nM via monitoring all mAb/dsRNA peaks as a cumulative population (Figure 3C). All models support that all binding sites on the dsRNA are equivalent and occur independent of the next mAb consistent with earlier reports (34). It’s notable that this MP *K*_D_ is an order of magnitude tighter than a BLI reported value (27-32 nM) that is likely influenced by avidity effects, a higher salt concentration (100 mM NaCl), or buffer excipients (e.g., tRNA, tween-20) required for BLI experiments (30, 35). Our observations of multiple binding sites on the 400 bp dsRNA is also consistent with previous AFM analysis, which reports how multiple J2 mAbs will bind a single dsRNA with a footprint of ∼40 bp (10), and is an example of the unique structural insights offered by MP. We then repeated the experiment in TE buffer with 150 mM NaCl to test whether increased amounts of the J2 mAb was able to promote formation of the mAb/dsRNA complex at high salt. Supplementary Figure S2 shows how the 1:1 complex remains inhibited up to 100 nM J2 mAb and supports the decision to proceed with lower ionic strength.

**Figure 3.**
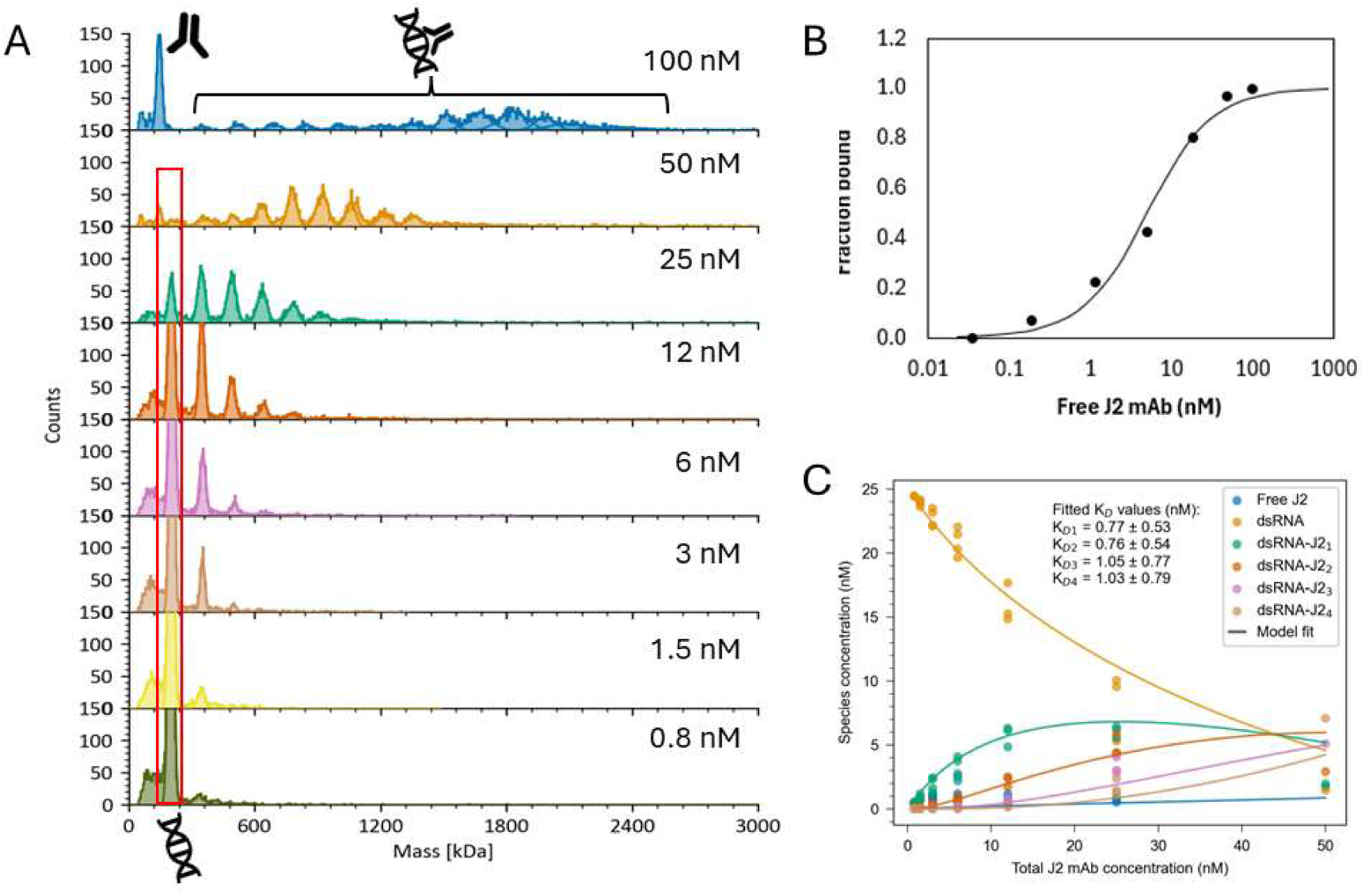
Binding affinity (K_D_) of the J2 mAb for dsRNA. (A) Mass photometry histograms of 2 nM dsRNA substrate with increasing amounts of the J2 mAb. The free J2 mAb is observed at 150 kDa, the free dsRNA is observed at 220 kDa, and mAb/dsRNA complexes are observed above 350 kDa. Data were prepared and collected in TE buffer supplemented w 50 mM NaCl. (B) Determination of the K_D_ between the J2 mAb and 400 bp dsRNA substrate. The fraction bound was determined using the Gaussian fits for the free dsRNA peak (denoted by the red box in Fig 3A). The K_D_ was then determined by the quadratic equation defined in the methods and materials. (C) Titration fits for binding affinity of J2 mAb to the 400 bp dsRNA substrate with 1-4 assumed binding sites per dsRNA molecule.

### mAb-observed detection of dsRNA on MGUC

The low molecular weight species observed in the dsRNA data on MGNA slides can obscure detection of small amounts of unbound mAb. Hence, we surmised that uncoated slides would simplify analysis by detecting only the free J2 mAb and the mAb/dsRNA complex while repelling free nucleotides. We prepared a mAb/dsRNA sample at a 2:1 ratio in TE buffer and analyzed the sample on either a MGNA (RNA-observed) or uncoated (mAb-observed) slide. Figure 4 shows how the MGNA slide presents peaks representing the free J2 mAb (146 kDa), the free dsRNA (206 kDa) and the mAb/dsRNA complex with one (352 kDa), two (500 kDa), or three (646 kDa) bound mAbs. It also shows how the uncoated slide only has binding peaks corresponding to the free J2 mAb (143 kDa), the three mAb/dsRNA complexes (i.e., 334 kDa, 484 kDa, and 627 kDa), and one unbinding peak that corresponds to the mass of the free dsRNA (188 kDa). The absence of the binding peak for dsRNA in the uncoated histogram supports that the unbinding peak represents dsRNA molecules that are released from the mAb after a binding event as a mAb/dsRNA complex. The relative counts for the unbinding dsRNA (4054 +/- 939 counts) are also within error of the summed binding peaks for the mAb/dsRNA complex (4021 +/- 1014 counts). This unbinding peak in the uncoated data is therefore a useful confirmation of the dsRNA species captured by the J2 mAb even when multiple mAbs are bound.

**Figure 4.**
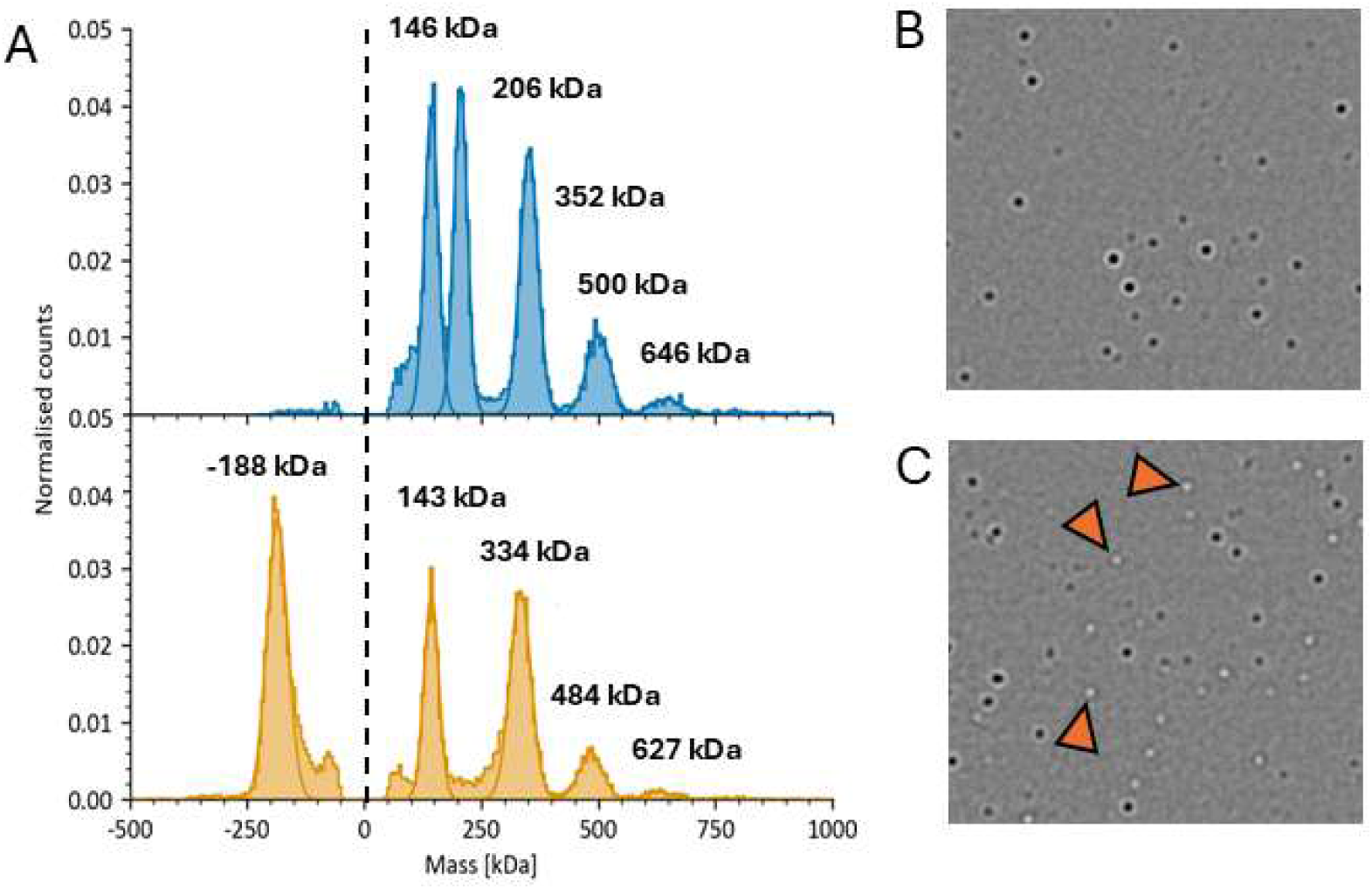
Detection of dsRNA species on cation-coated (blue) and uncoated (yellow) slides. (A) Mass photometry histograms of the mAb/dsRNA complex on MG-NA (*top*) or MG-UC (bottom). Samples were prepared in TE buffer with 50 mM NaCl at a 2:1 ratio for the J2 mAb and 400 bp dsRNA reagents. The dotted line at the origin is a visual aid for separating the positive binding masses from the negative unbinding masses. (B) Representative field of view image of binding events observed on MG-NA slides corresponding to Figure 4A. (C) Representative field of view image of binding events observed on MG-UC slides corresponding to Figure 4A. Examples of unbinding events are denoted by orange triangles.

A MassFluidix experiment was performed to test whether a liberated dsRNA is released by the J2 mAb and will rebind the surface as new binding event if any mAbs remain on the dsRNA. In such a scenario, we expect a different relative histogram distribution between static and MassFluidix experiments. MassFluidix experiments are conventionally done to study weakly interreacting complexes by recording MP data of rapidly diluted samples (36, 37); however, the continuous flow of the system selectively isolates molecules approaching the focal point, ensuring that molecules unbinding from the surface are washed away rather than being recounted during the experiment. Supplementary Figure S3 compares the MP histograms of the mAb/dsRNA sample on an uncoated slide from a standard equilibrium experiment and a MassFluidix chip. There are two takeaways from this experiment. First, both the static and MassFluidix experiments show very good agreement between the observed mAb/dsRNA complexes, with only one observed species (465 kDa) not aligning with a well-defined multiple of mAb bound to the dsRNA substrate. Instead, this 433 kDa peak is likely an artefactual amalgam of complexes with 1 or 2 bound mAbs per dsRNA merged into a single broad distribution. The presence of higher-order species (i.e., 1281 and 1440 kDa) in the MassFluidix data result from the higher preparative concentration of the sample (∼125-fold) not achieving equilibrium after rapid dilution. All species, however, follow the additive 150 kDa pattern that supports multiple mAbs bound to the dsRNA. The second takeaway is that both experiments show a single unbinding peak that represents the liberated dsRNA.

### Characterization of dsRNA in commercial mRNA

We next applied the learnings for MP detection of dsRNA to a commercial mRNA sample. ELISA analysis of the GFP mRNA sample benchmarks the dsRNA content at 0.11% (Figure 5) (8), which means a mRNA concentration of >100 nM is required to detect dsRNA within the sample. MP experiments are typically unable to be done at such high concentrations because too many events will saturate the field of view and obscure individual binding events. Uncoated slides, however, will only register mAb and mAb-bound RNA species leaving the bulk RNA populations undetected. To this end, we developed an assay using a fixed amount of J2 mAb (25 nM) to ensure the system remained above the K_D_ while defining the upper limit for experimental binding counts. This configuration ensures that only mAb-bound complexes are detected, as the RNA analyte does not interact with the uncoated slide surface.

**Figure 5.**
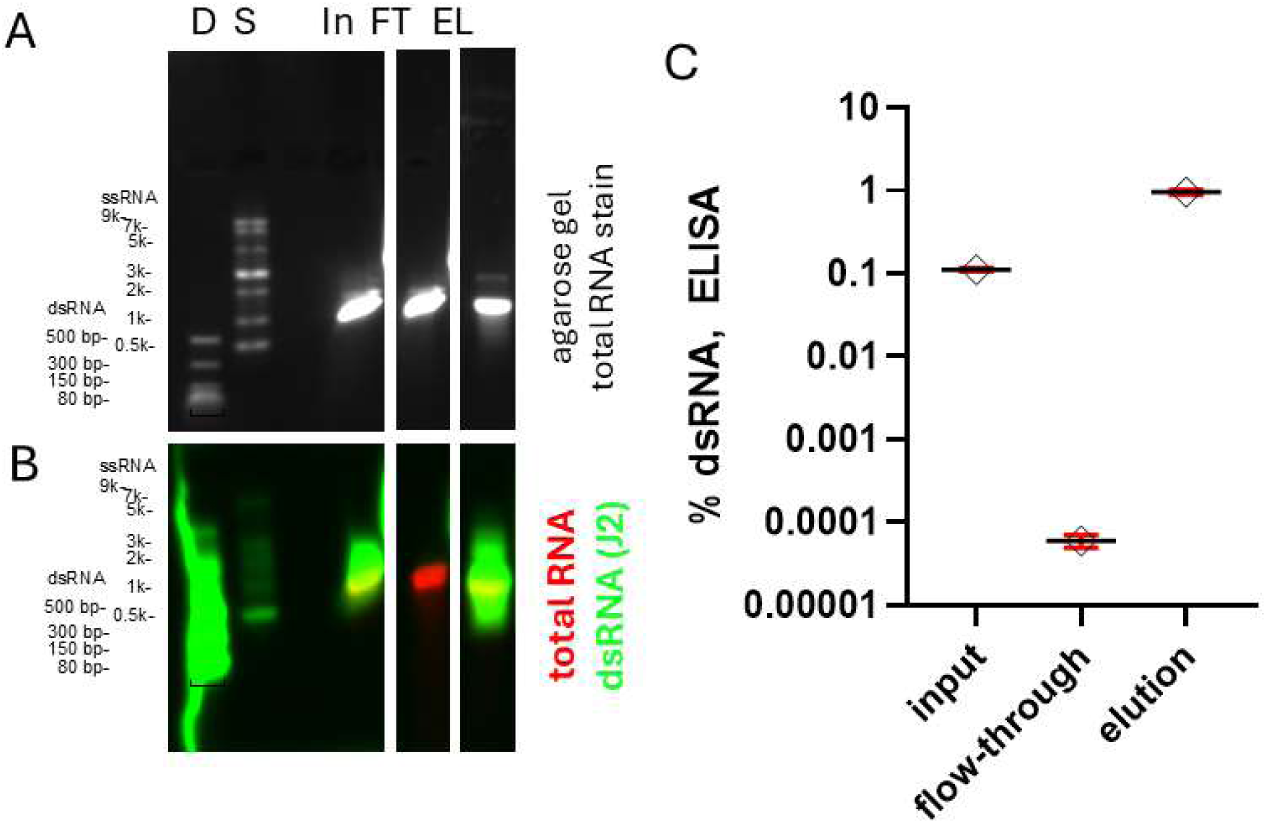
Analysis of affinity-purification samples for uncapped GFP mRNA (A) Agarose gel of mRNA samples stained for total-RNA. Lanes are defined as: D; dsRNA ladder, S, single-stranded RNA ladder, In, input mRNA; FT, Flow-Through fraction; EL, Elution fraction. (B) Dual color immuno-northern blot of the agarose gel presenting the total RNA stain (red) overlaid with the J2-chemiluminescent signal (green). (C) Quantification of dsRNA levels in the uncapped GFP fractions using the Vazyme dsRNA ELISA kit. The dsRNA content of each fraction ([µg dsRNA/µg total RNA]*100) is plotted, with error bars representing the standard deviation from replicate measurements.

Figure 6 presents two experiments to demonstrate the necessity and performance of uncoated slides for detecting and characterizing dsRNA impurities in an mRNA sample by MP. The first experiment validates the selectivity of the J2 mAb for dsRNA by incubating 25 nM J2 mAb with 5 nM GFP mRNA and collecting MP measurements on catatonically coated MG-NA slides to observe all RNA events (Figure 6A, top). Note that 5 nM RNA was used to keep the number of observed events from saturating the instrument detector. The resulting histogram showed three peaks that are attributed to the J2 mAb (150 kDa), a peak for the intact monomer (231 kDa), and a small peak for the RNA multimer (463 kDa). Note that the two RNA peaks are consistent with the RNA species observed without the J2 mAb in Figure 1A and therefore neither represent a mAb/dsRNA complex. The 463 kDa species is particularly interesting as the observed mass supports the presence of a larger RNA (presumably a multimer) that is undetectable by the J2 mAb as there is no +150 kDa species in this histogram relative to the profile in Figure 1A. Rather, the more likely scenario is that this 463 kDa species represents a transient dsRNA multimer or aggregate as described by Webb et al. (21) and no significant dsRNA populations are detected in this scenario.

**Figure 6.**
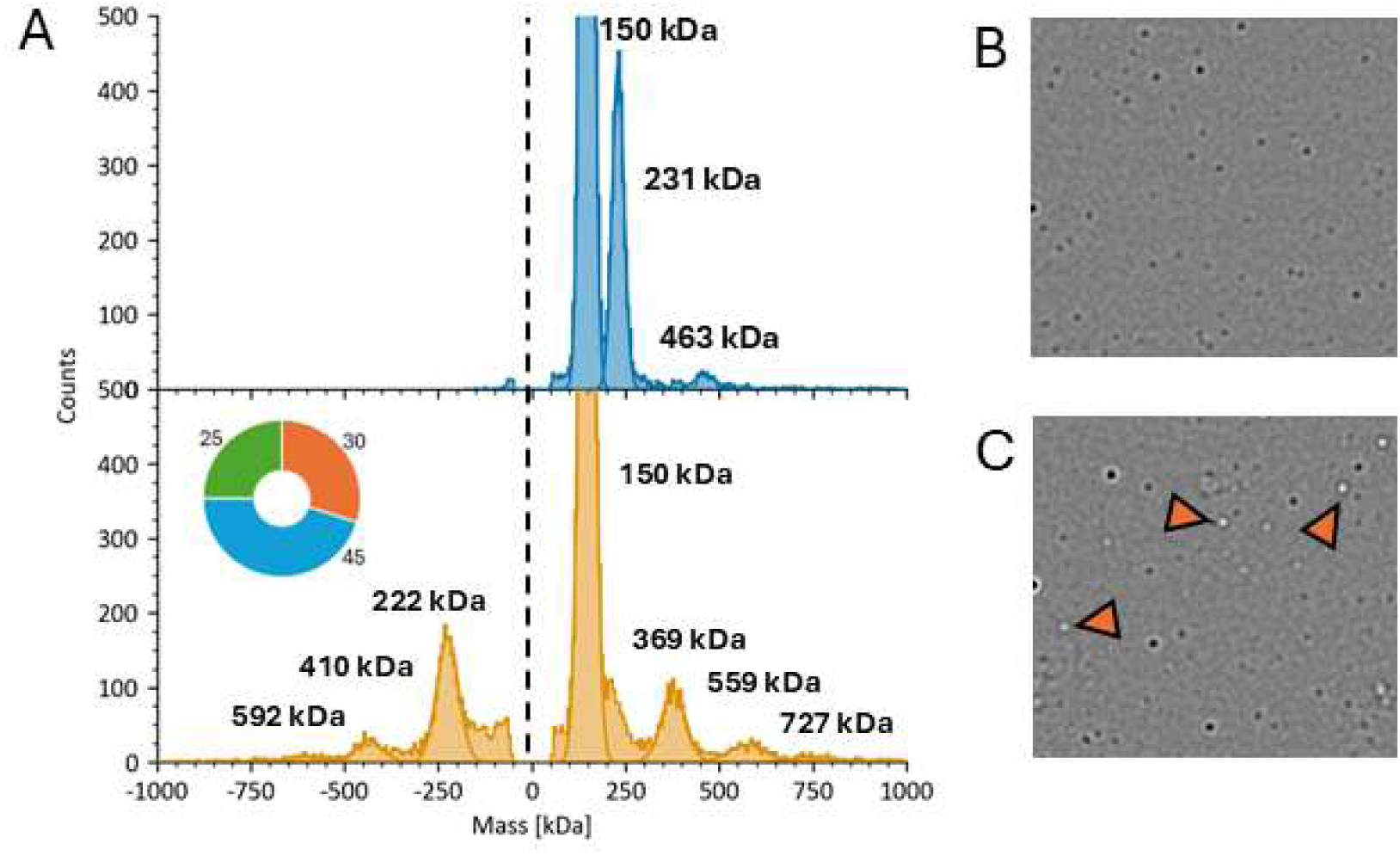
Detection of dsRNA species in an mRNA sample on cation-coated (blue) and uncoated (yellow) slides. (A) Mass photometry histograms of the mAb/dsRNA complex on MG-NA (*top*) or MG-UC (bottom). Samples for analysis on MG-NA were prepared in by mixing J2 mAb (25 nM) and GFP mRNA (5 nM). Samples for analysis on MG-UC were prepared in by mixing J2 mAb (25 nM) and GFP mRNA (120 nM) in TE buffer with 50 mM NaCl. The dotted line at the origin is a visual aid for separating the positive binding masses from the negative unbinding masses. (B) Representative field of view image of binding events observed on MG-NA slides corresponding to Figure 5A. (C) Representative field of view image of binding events observed on MG-UC slides corresponding to Figure 5A. Examples of unbinding events are denoted by orange triangles.

The second experiment uses a higher mRNA concentration (120 nM) for the assay on uncoated slides to illustrate a heterogeneous dsRNA profile in the GFP mRNA (Figure 6A, bottom). Recall the uncoated slides allow us to use a higher concentration of RNA, which will not bind the surface without a the J2 mAb, to detect the rare dsRNA impurity as a mAb/dsRNA complex. The resulting histogram revealed four binding populations (150 kDa, 369 kDa, 559 kDa, and 727 kDa) and three unbinding populations (222 kDa, 410 kDa, and 592 kDa). The net difference of the masses for the unbinding events are ∼150 kDa lower than the masses observed for the binding events and support that these are *bona fide* dsRNA species observed on the uncoated slide via the J2 mAb. Moreover, it is notable that these are three distinct species as the three binding peaks correlate to the three unbinding peaks by a net mass difference of 150 kDa (1x J2 mAb) and do not collapse into a single dsRNA population as is observed for the dsRNA standard (Figure 4). The unbinding 222 kDa species indicates that there is some population of RNA consistent with the mass of an intact monomer that is recognized by the J2 mAb, either via a short annealed abortive RNA or a long hairpin ≥40 bp. The unbinding 410 kDa and 592 kDa species indicate the presence of distinct aggregate or multimer species within the sample. A broad shoulder to the right of the unbinding 222 kDa species is also notable as it likely represents a significant presence of poorly defined low mass species from abortive transcripts and degradation products.

Constant-flow MassFluidix analysis on uncoated flow chips was next used to corroborate the presence of dsRNA impurities in the mRNA sample and eliminate the possibility of experimental bias. This dynamic measurement ensures that the recorded events reflect the true sample composition rather than localized rebinding of dissociated particles over time. Because the relative populations aligned with previous static MP observations, these results confirm the robustness of the assay for mRNA purity characterization. Supplementary Figure S4 shows the MP histograms for both experiments with a major binding peak for the free J2 mAb (150 kDa) and several minor peaks representing an array of mAb/dsRNA complexes (375 kDa, 552 kDa, 727 kDa). Three unbinding peaks are observed at 220 kDa 400 kDa, and 600 kDa, each having a differential mass of ∼150 kDa from the corresponding mAb/dsRNA species. The corresponding MFx experiment again corroborated the results of the static MP experiment.

### Characterization of dsRNA in processed mRNA fractions

We next processed the GFP mRNA with dsRNA affinity resin and collected the Flow Through and Elution fractions to probe dsRNA impurities with the J2 mAb as described above. An initial mRNA profile was collected on MG-NA slides to establish the “total RNA” profile of both fractions (Figure 7A). The Flow Through fraction shows a very similar profile to the unprocessed mRNA (Figure 1A), with a dominant peak for the intact monomer (234 kDa), a small population of the RNA multimer (468 kDa), and a low population of broadly distributed low molecular weight impurities to the left of the intact monomer. This result is consistent with the dsRNA content of only 0.1% of total RNA by ELISA (Figure 5C). The Elution fraction shows a slightly different profile, whereby the most prevalent difference from the Flow Through fraction is a significantly increased amount of low molecular weight species relative to the 232 kDa species and a diminished amount of the RNA multimer (459 kDa). We interpret this difference to mean that the dsRNA affinity column has removed both a significant amount of low molecular weight dsRNAs from the mRNA sample and an mRNA species with near the intact mass (230 kDa) with dsRNA character different from what is in the Flow Through fraction. We cannot clearly define the nature of the 232 kDa species, but rationalize the presence of either the intact mRNA monomer hybridized with a short abortive RNA sequence or a complex hairpin structure that was recognized by the column. Moreover, the diminished amount of multimer species in the Elution fraction is interesting because it indicates that this species is not a dsRNA species recognized by the dsRNA affinity resin. Hence, it is likely that the multimer species observed in the GFP Input and Flow Through fractions are transient, concentration-dependent species demonstrated by Webb et al. (21). The Elution fraction therefore contains RNAs that are less likely to form these multimer species.

**Figure 7.**
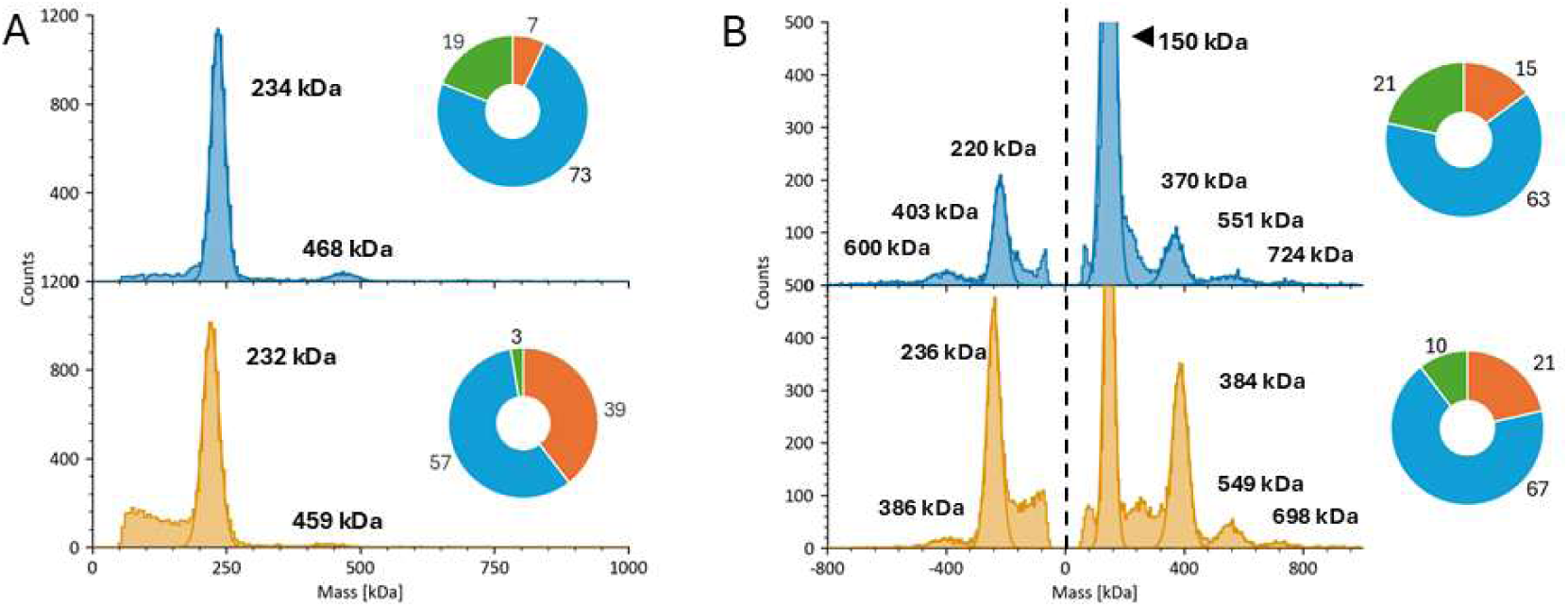
Mass photometry histograms of the flow through (blue) and elution (yellow) fractions of GFP mRNA after dsRNA affinity chromatography. (A) Mass photometry histograms of GFP fractions (5 nM) on MG-NA slides prepared in IDTE buffer with 50 mM NaCl. The inset pie chart represents the relative amount of low mass (orange), monomer (blue) and aggregate (green) species determined by percentage of counts in gated MP data. (B) Mass photometry histograms of GFP fractions (5 nM) with 25 nM J2 mAb. Samples were prepared in IDTE buffer with 50 mM NaCl and measured on MG-UC slides. The pie charts represent the relative amount of species observed in the unbinding data and are presented as defined in Fig 7A.

A second experiment was then performed with the addition of the J2 mAb on uncoated slides to establish the profile of dsRNA impurities within these mRNA fractions (Figure 7B). The MP histograms for the Flow Through and Elution fractions show the presence of dsRNA species being near the mass of an mRNA monomer (i.e., 220-236 kDa). This dominant dsRNA species is interesting because we can resolve a small but meaningful mass difference is between the Flow Through and Elution fractions in both the binding and unbinding data (Table 2). Separation of distinct populations by the column resin is supported by reduction of the standard deviation of the input fraction relative to both the Flow Through and Elution fractions. This reduction in σ (peak width) is also observed for each measured peak and indicates a more homogeneous population relative to the Input fraction. While we cannot specifically identify the nature of dsRNA present in these fractions (e.g., 3’ loopback or hybridization between abortive transcripts), the observed mass difference indicates that the column is removing a distinct dsRNA that may be shorter than the 30-40 bp length recognized by the J2 mAb. It is also notable that the observed aggregate masses are not true multiples of the mRNA monomer and indicate they may be sequence-dependent annealing between shorter truncated RNAs.

**Table 2.** Observed mass and sigma for the monomer-like peak observed within GFP mRNA fractions separated on a dsRNA affinity column. Data represent the average of three measurements performed on uncoated slides (Figure 7B).

|  |  | Input | Flow Through | Elution |
| --- | --- | --- | --- | --- |
| Binding | Counts | 1,235 ± 301 | 1,070 ± 363 | 3,440 ± 536 |
|  | Mass (kDa) | 369 ± 11 | 370 ± 3 | 384 ± 5 |
|  | σ (kDa) | 36 ± 4 | 30 ± 3 | 27 ± 1 |
| Unbinding | Counts | 1,543 ± 436 | 1,589 ± 607 | 3,845 ± 411 |
|  | Mass (kDa) | 222 ± 9 | 220 ± 3 | 236 ± 4 |
|  | σ (kDa) | 26 ± 1 | 22 ± 3 | 24 ± 2 |

We next divided the MP data for each mRNA fraction into three gates to analyze the relative populations of low mass species (50-150 kDa), the species that aligns with the intact monomer (150-275 kDa), and larger aggregate/multimer species (>275 kDa). Figure 8A compares the distributions of total RNA of each mRNA fraction from measurements on MG-NA without J2 mAb. The gated analysis further quantifies how the Elution fraction is significantly enriched for low mass RNAs relative to depleted monomer and multimer species (Figures 6 and 7). Figure 8B compares the dsRNA impurities on uncoated slides for the three mRNA fractions. The Elution fraction again shows a decrease in the relative amount of larger aggregate/multimer RNAs with an increase in the low mass and monomer-like populations relative to the Flow Through. Both the Flow Through and Elution fractions exhibit a different dsRNA profile from the Input fraction, which we interpret to mean that the process of removing dsRNA via the affinity column may in itself disrupt some weakly hybridized transient species reported by Webb et al (21) to enable the separation of distinct species observed in Figure 7 and Table 2. Overall, MP detects a significant amount of low mass dsRNAs from the GFP elution sample. This observation is enabled by the concentration of dsRNAs on the affinity columns, as these small dsRNAs are not detectible in either the input or flow through fractions due to the low titers (Figure 5C). The elution fraction was also enriched in high-molecular weight dsRNAs such as those at 384 and 549 kDa. A dsRNA detection reagent that recognizes shorter dsRNA segments would likely give better resolution of these populations and is an exciting opportunity for further development of dsRNA characterization by MP.

**Figure 8.**
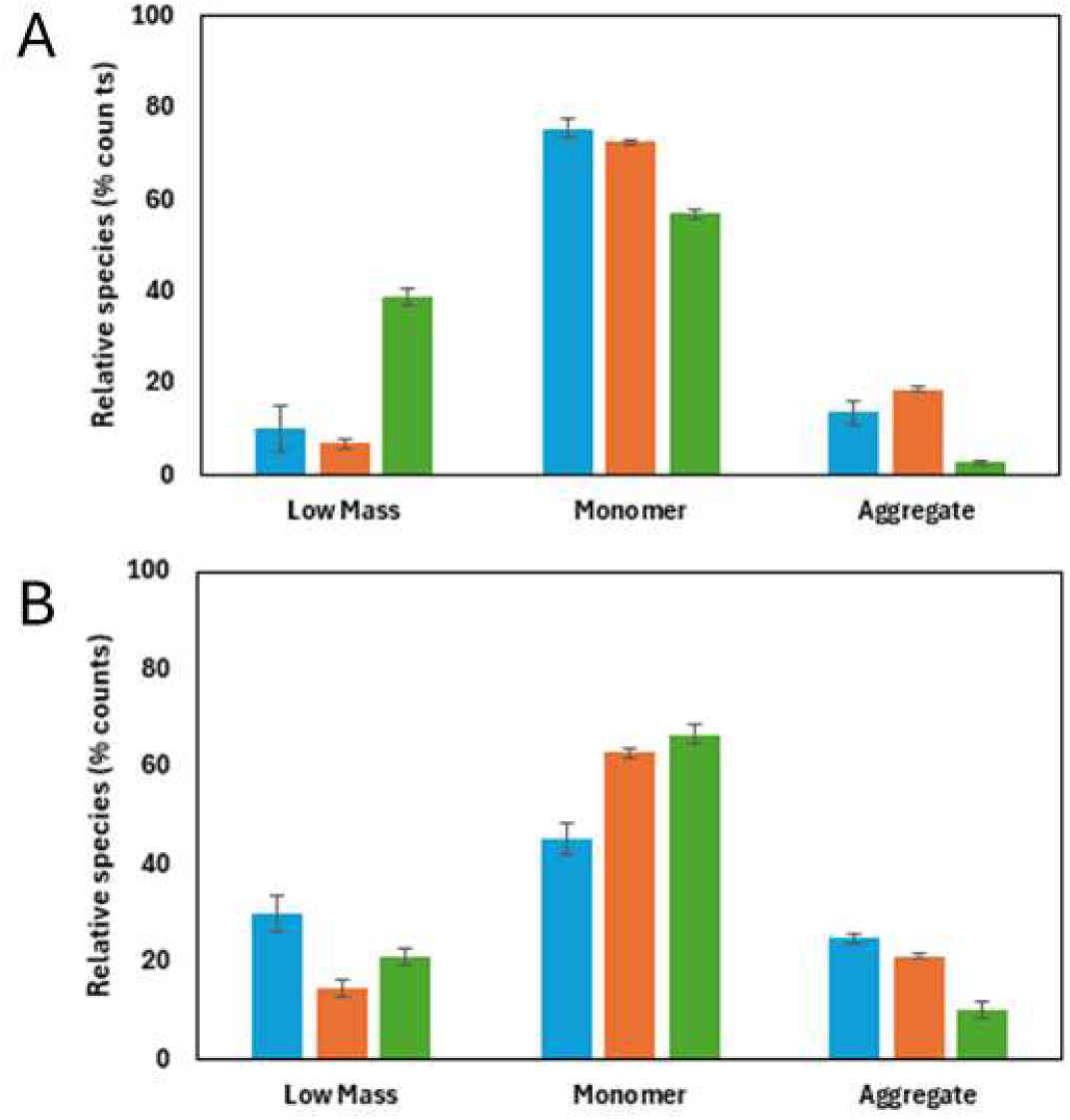
RNA content for input (blue), flow through (orange), and elution (green) fractions of GFP mRNA from purification on dsRNA affinity resin. (A) Total RNA species from MP data collected on MG-NA and gated for low mass species (0-200 kDa), intact monomer (200-300 kDa), and aggregate species (300+ kDa). (B) dsRNA species from unbinding MP data collected on MG-UC. Data were gated as described in Fig 8A.

### Quantification of %dsRNA

Quantification of the relative %dsRNA can be calculated from the MP data. Specifically, the relative amount of dsRNA impurity can be determined as the ratio of dsRNA unbinding counts from uncoated slides over the total counts from MGNA after all counts are corrected for mass-weighting and concentration. Table 3 presents the MP-determined amounts of dsRNA impurities by gated populations or the summed total. Note that we have chosen to treat the degradation products and higher order species (aggregates) as bulk interval regions relative to a “monomer-like” species (based on the observed mass) to simplify the characterization in this work; however, one could subdivide each fraction further if they desired. For the analysis presented herein, we focus on the total %dsRNA levels for comparison between the fractions because these are comparable to the bulk analysis provided by the immunoblot and ELISA assays. Comparison of the %dsRNA levels for the three fractions shows a systematic decrease between the Input and Flow Through fractions followed by a significant increase in %dsRNA levels for the Elution fraction.

**Table 3.**
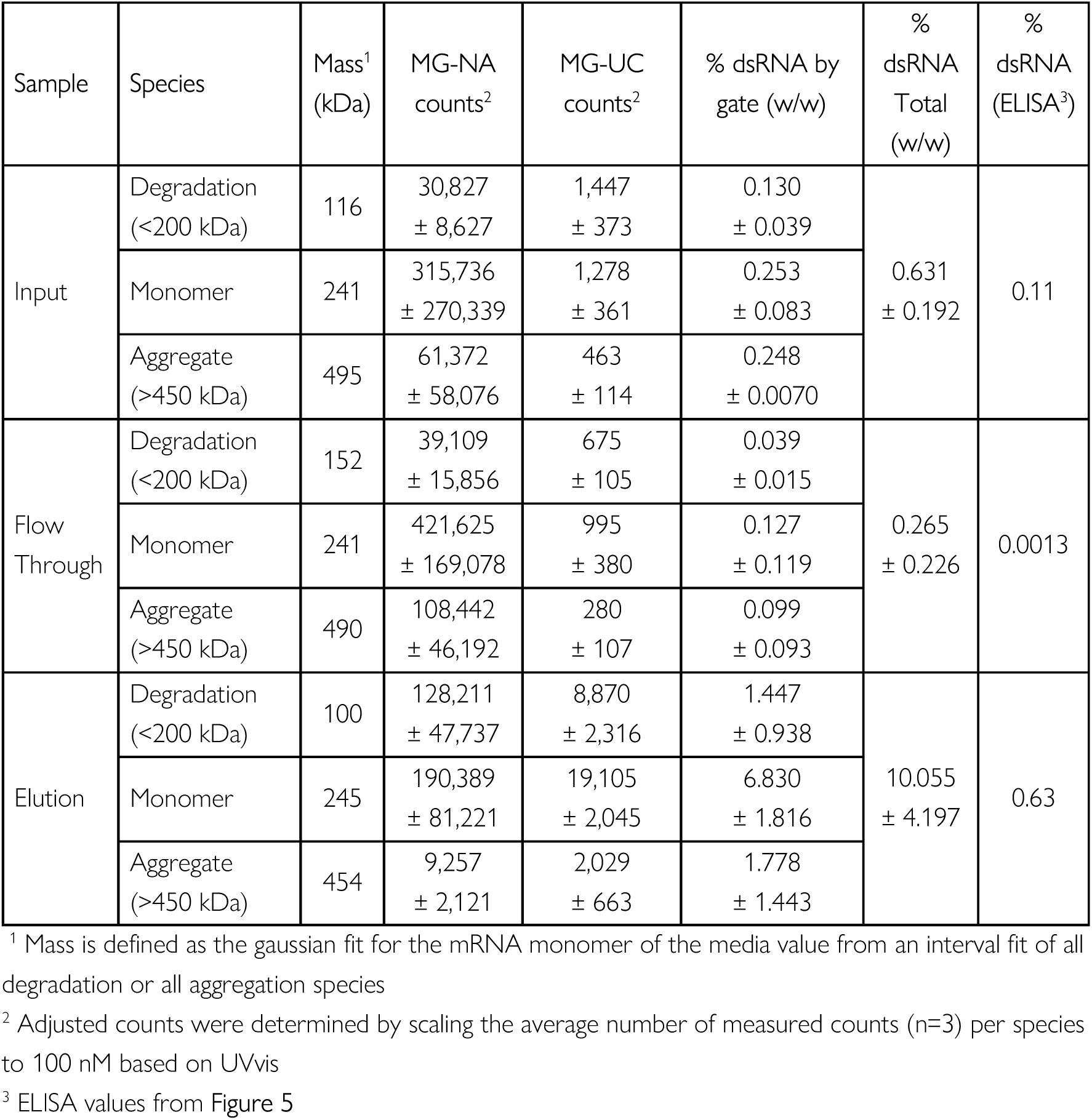
Estimation of %dsRNA impurities from MP data of the column fractions collected on MG-UC and MG-NA (Figures 6 and 7). See methods section for experimental details.

| Sample | Species | Mass<sup>1</sup><br>(kDa) | MG-NA<br>counts<sup>2</sup> | MG-UC<br>counts<sup>2</sup> | % dsRNA by<br>gate (w/w) | %<br>dsRNA<br>Total<br>(w/w) | %<br>dsRNA<br>(ELISA<sup>3</sup>) |
| --- | --- | --- | --- | --- | --- | --- | --- |
| Input | Degradation<br>(<200 kDa) | 116 | 30,827<br>± 8,627 | 1,447<br>± 373 | 0.130<br>± 0.039 | 0.631<br>± 0.192 | 0.11 |
|  | Monomer | 241 | 315,736<br>± 270,339 | 1,278<br>± 361 | 0.253<br>± 0.083 |  |  |
|  | Aggregate<br>(>450 kDa) | 495 | 61,372<br>± 58,076 | 463<br>± 114 | 0.248<br>± 0.0070 |  |  |
| Flow<br>Through | Degradation<br>(<200 kDa) | 152 | 39,109<br>± 15,856 | 675<br>± 105 | 0.039<br>± 0.015 | 0.265<br>± 0.226 | 0.0013 |
|  | Monomer | 241 | 421,625<br>± 169,078 | 995<br>± 380 | 0.127<br>± 0.119 |  |  |
|  | Aggregate<br>(>450 kDa) | 490 | 108,442<br>± 46,192 | 280<br>± 107 | 0.099<br>± 0.093 |  |  |
| Elution | Degradation<br>(<200 kDa) | 100 | 128,211<br>± 47,737 | 8,870<br>± 2,316 | 1.447<br>± 0.938 | 10.055<br>± 4.197 | 0.63 |
|  | Monomer | 245 | 190,389<br>± 81,221 | 19,105<br>± 2,045 | 6.830<br>± 1.816 |  |  |
|  | Aggregate<br>(>450 kDa) | 454 | 9,257<br>± 2,121 | 2,029<br>± 663 | 1.778<br>± 1.443 |  |  |

<sup>1</sup> Mass is defined as the gaussian fit for the mRNA monomer of the media value from an interval fit of all degradation or all aggregation species
<sup>2</sup> Adjusted counts were determined by scaling the average number of measured counts (n=3) per species to 100 nM based on UVvis
<sup>3</sup> ELISA values from **Figure 5**

The MP values in Table 3 are collectively higher than those determined by ELISA (Fig 5) and led us to consider the experimental factors driving either analysis. Firstly, the two methods specifically use different antibodies for analysis, where we used the J2 mAb and the ELISA relies on a proprietary set of antibodies (i.e., M2 and M5). Characterization of these reagents shows enough differences in affinity, binding orientation of the substrate, and minimal dsRNA footprint (27 v >40 bp) to suggest different binding modes of either reagent (30, 38). Second, the ELISA kit use proprietary buffers for the antibody incubation and wash protocol. A survey of commercial ELISA kits indicate the widespread use of phosphate or Tris buffered saline, both of which have a higher ionic strength (IC ∼ 170 mM) than the buffer developed for our MP work (I_C_ ∼ 55 mM). We therefore chose to bridge the two assays using a J2-based immunoblot to compare salt conditions with a single antibody for detection. Figure 9 presents the immunoblot performed at either standard (150 mM) or reduced (50 mM) salt concentrations and shows a significant increase in the signal strength at lower salt relative to standard conditions with a *p-value* of 0.00132. These results agree with the MP data showing salt sensitivity of the mAb/dsRNA complex (Fig 2) and J2 titration data at 50 mM (Fig 3) or 150 mM (Supplementary Figure 2) NaCl.

**Figure 9.**
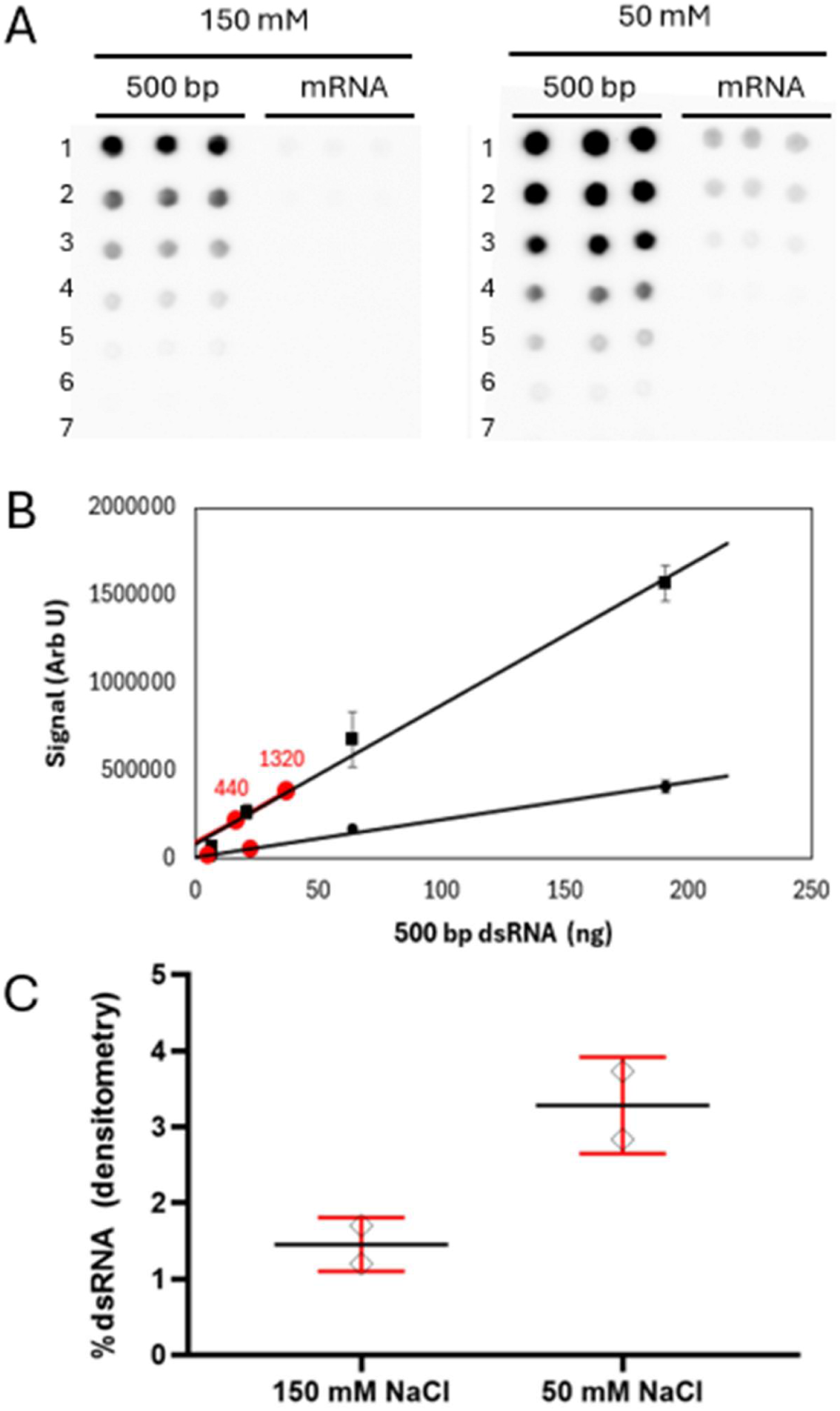
Immunodetection of dsRNA impurities in GFP mRNA with varied salt concentrations. (A) Representative dot blot images comparing target detection and assay sensitivity in 150 mM (left) and 50 mM (right) buffer conditions. A known 500 bp double-stranded standard and a GFP mRNA sample were spotted in triplicate to evaluate relative impurity content. Samples were applied using a 7-point dilution series (rows 1–7, top to bottom). Initial loading concentrations were 820 ng/µL for the 500 bp dsRNA standard and 660 ng/µL for the GFP mRNA. (B) Standard curve of the 500 bp dsRNA standard performed in 50 mM (square) or 150 mM (circle) buffer conditions. The fit values for two mRNA samples (440 and 1320 ng) is plotted as red circles on the standard curves of either buffer condition. (C) Quantification of dsRNA levels using aJ2-HRP mAb. The dsRNA content of each fraction ([µg dsRNA/µg total RNA]*100) is plotted, with error bars representing the standard deviation from replicate measurements.

## DISCUSSION

Current analytical technologies face persistent challenges in the detection and characterization of dsRNA impurities in mRNA therapeutics. For example, immuno-based methods like ELISA and dot blot return a single, bulk %dsRNA value per sample, offering no insight into the underlying subpopulation distribution. This limits the utility of these methods for understanding dsRNA impurities and providing little actionable guidance for process optimization or root-cause investigation. The work presented in this study demonstrates how MP addresses this critical unmet need in the detection and semi-quantitative profiling of dsRNA containing impurities in mRNA preparations. MP provides single-molecule resolution of the intact RNA species, enabling direct measurements of heterogeneous RNA populations within complex samples. In contrast to enzymatic, chromatographic, or sequencing-based assays, MP resolves and quantifies distinct populations of RNAs with dsRNA features without amplification or extensive sample processing (Figure 1 and 2).

The J2 mAb is a highly selective reagent for A-form dsRNA structures and widely used across immuno-based assays; however, few biophysical studies have been performed to define the nature of a stable mAb/dsRNA complex. While two BLI-based studies have noted the relative salt-sensitivity of the interaction (30, 35), neither fully explores the salt-dependence of the complex. We therefore built our MP-based assay from a salt titration that showed the mAb/dsRNA complex is best represented in TE with 50 mM salt (Figure 2). A K_D_ of ∼1 nM was determined for the J2 mAb under these conditions (Figure 3). This value is significantly tighter than the 27 nM value from BLI (35), and agrees with the salt dependence of the mAb/dsRNA complex observed in Figure 2B. Furthermore, researchers should take these buffer and salt effects into account for other immunodetection methods that may alter quantification of the dsRNA impurities in their drug substance.

We next developed an MP assay on uncoated slides for a mAb-observed detection for two reasons. First, nucleotides will not interact with an uncoated surface (18) and allows selective control of the relative counts in the assay with a fixed amount of mAb (25 nM). All observed binding events on uncoated slides are therefore either the free J2 mAb or a mAb/dsRNA complex (Figure 4). This approach represents a paradigm shift in impurity profiling by MP, where surface-level repulsion of the bulk analyte (e.g., RNA on uncoated slides) is harnessed to overcome the dynamic concentration range limitations of a typical experiment. This ’selective capture’ design enables the use of higher sample concentrations to facilitate sensitive detection of lowly populated impurities with single-molecule precision that would otherwise be obscured by the primary species. Second, the strong repulsion of the uncoated slide surface with the RNA generates an unbinding peak represents a free RNA containing a dsRNA species after it is liberated from the complex following mAb binding. These unbinding events act as both an internal confirmation for binding events and fingerprinting of the dsRNA and aggregate species by relative mass. The unbinding peaks are also important because they represent the cumulative peak that may result from multiple dsRNA features (i.e., multiple bound mAbs). However, due to the concentration dependent self-association of RNA (21), it is possible that some of the aggregates exhibit dsRNA character in the observed unbinding phenotype. Hence, it is likely that RNA aggregates are present in the unbinding MP data, especially considering that the unbinding signals were similar in the input, flow through, and elution fractions despite orders-of-magnitude differences in dsRNA content (i.e., 0.1%, 0.0006%, and 1% respectively) (Figure 5C).

We used GFP mRNA to demonstrate how the MP-based method translates from a dsRNA standard to mRNA fractions processed with a dsRNA affinity resin (Figures 6 and 7). Several dsRNA species are identified by the J2 mAb to have masses consistent with the mRNA monomer (220-230 kDa) and larger aggregate species (∼400 and 600 kDa). The elution fraction is highly enriched with small species that could represent small dsRNA hybrids or degradation products. The larger dsRNA species reported by others (7, 8) were not as robustly detected in these MP experiments (Figure 8). We hypothesize that these larger species may represent transient high-molecular weight RNAs identified by the J2 mAb consistent with the concentration-dependent RNAs reported by Webb et al (21). Regardless of their nature, the presence of high-molecular weight RNA multimers in the flow-through fractions but not the elution is evidence that the dsRNA-specific affinity ligand faithfully discriminates high molecular weight RNA multimers from true dsRNA molecules.

Quantification of dsRNA impurities is commonly performed using ELISA and immunoblot methods, both of which are listed by the US Pharmacopeia (39) and well represented in the field (8, 40, 41). To cross-validate the salt sensitivity of the J2 mAb observed by MP, we benchmarked it against a J2-HRP–based dot blot as an orthogonal method. Consistent with the MP-derived titration and fractionation data, the dot blot showed significantly stronger signal under reduced-salt conditions (50 mM) than standard conditions (150 mM NaCl; *p* = 0.00132) (Fig 9), confirming that the salt-dependent binding behavior of J2 is intrinsic to the antibody and not an artifact of the MP platform. This agreement, using the same antibody across both methods, also helps explain why the %dsRNA values obtained by MP exceeded those from ELISA, since the commercial ELISA kit relies on a distinct antibody pair (M2/M5) with different affinity, binding orientation, and dsRNA footprint than J2. Taken together, the %dsRNA values obtained across all three methods (Table 4) underscore that absolute dsRNA quantification remains highly method- and reagent-dependent, ranging from 0.11% by ELISA to 3.3% by J2-HRP dot blot under reduced-salt conditions, with MP (0.78%) falling within this range. As MP-based workflows continue to mature, we anticipate this single-molecule approach will offer increasingly robust, antibody-independent alternative for resolving these discrepancies and advancing standardized dsRNA impurity analytics in the field.

**Table 4.** Method comparison for dsRNA quantification. Single-molecule mass photometry analysis of molecular weight deviations. Relative error percentages provide a benchmark of measurement accuracy compared to sequence-derived values. A clear reduction in mass error is observed for higher-order nucleoprotein complexes.

| Method | Salt | Detection antibody | %dsRNA |
| --- | --- | --- | --- |
| ELISA | Proprietary | M5 | 0.11 +/- 0.006% |
| Dot blot | 150 mM NaCl | J2-HRP | 1.4 +/- 0.3 |
|  | 50 mM NaCl | J2-HRP | 3.3 +/- 0.6 |
| Mass photometry | 50 mM NaCl | J2 | 0.6 +/- 0.2 |

## CONCLUSIONS

The mRNA manufacturing field currently lacks a tool for understanding dsRNA impurities and informing process and product development. Bulk immuno-detection methods like ELISA and dot blot return only a single composite %dsRNA value, which informs teams that impurities exist without revealing which dsRNA containing species are present or where in the manufacturing process they arise. This study demonstrates that MP closes that gap by rapidly profiling distinct dsRNA containing populations within a single, minimally processed measurement. Such data give manufacturing teams a direct window into impurity composition rather than a single opaque number. Benchmarking against ELISA and dot blot further revealed that absolute %dsRNA values are highly method and reagent dependent, varying nearly an order of magnitude across current industry standard assays. This variability points to a broader problem, since without a shared reference standard, teams are often optimizing processes and setting specifications against numbers that are not directly comparable across methods, platforms, or sites. By resolving the specific molecular species behind a given %dsRNA result, MP offers manufacturing and CMC teams a path toward genuine root cause investigation and process optimization rather than pass or fail testing alone. This positions single molecule profiling as a practical tool for advancing dsRNA impurity control as mRNA therapeutics move toward more structurally complex, next generation constructs.

## Supporting information

Supporting images for the manuscript

## ACKNOWLEDGEMENTS

The authors thank Josh Bishop and Justin Benesch for their insightful discussions and critical reading of the manuscript. The authors also thank Catie Lichten for her support preparing the manuscript.

## AUTHOR CONTRIBUTIONS

M.J.R. performed all mass photometry experiments presented in this work. N.E.C. performed the purification of GFP mRNA fractions on dsRNA affinity resin and %dsRNA analysis via ELISA and dot blot. K.F. and M.J.R. performed the MassFluidix experiments on dsRNA complexes. A.R.O. performed data analysis related to binding affinity models. M.J.R., N.M., and G.K. contributed to the semi-quantification analysis of dsRNA by mass photometry. M.J.R. and S.C. conceptualized experiments and wrote the manuscript. All authors reviewed, contributed to, and approved the final manuscript.

## SUPPLEMENTARY DATA

Supplementary Data are available at NAR online.

## CONFLICT OF INTEREST

M.J.R., K.F., S.C., N.M, and G.K. are employees of Refeyn, Ltd. N.E.K. is an employee of Repligen. A.R.O is a consultant for Refeyn Ltd. Refeyn Ltd. manufactures mass photometer instruments. Repligen manufactures and sells bioprocessing materials and related products.

## DATA AVAILABILITY

Datasets generated and analyzed in the current study are available from the corresponding authors on reasonable request.

## Notes

### Competing Interest Statement

M.J.R., K.F., N.M, S.C. and G.K. are employees of Refeyn, Ltd. C.N.K. is an employee of Repligen. A.R.O is a consultant for Refeyn Ltd. Refeyn Ltd. manufactures mass photometer instruments. Repligen manufactures and sells bioprocessing materials and related products.

### Summary of Updates

Added section on J2 quantification by dot blot and MP for comparison against the ELISA data

