## Supporting images for the manuscript for "Double-Stranded RNA Profiling with Mass Photometry"

Supplementary Figure S1. Effect of ionic strength on protein and dsRNA calibration curves.

Supplementary Table S1. Fits for the calibration curves presented in Supplementary Figure 2

Supplementary Figure S2. Comparison of static and MassFluidix MP data for the mAb/dsRNA complex using the 400 bp dsRNA standard w TandemMP analysis

Supplementary Figure S3. Comparison of static and MassFluidix MP data for the mAb/dsRNA complex using GFP mRNA w TandemMP analysis

Supplementary Figure S4. Mass photometry GFP mRNA on coated and uncoated slides.

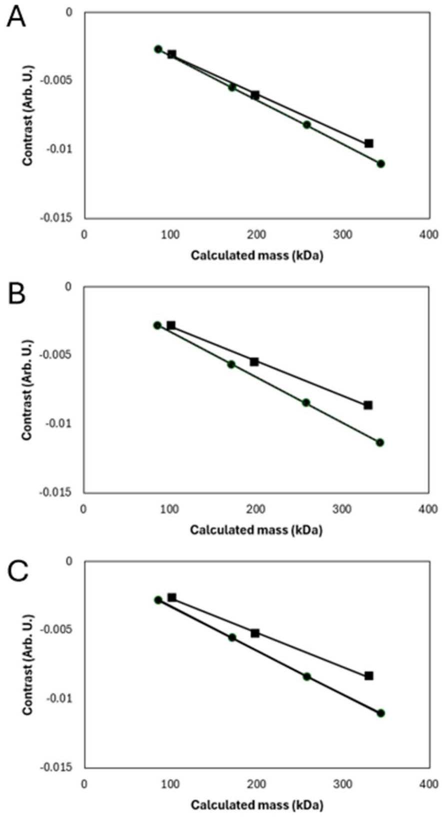


Supplementary Figure S1. Effect of ionic strength on calibration curve for protein (circles) and dsRNA (squares). Calibration samples were prepared in TE buffer with (A) no added salt, (B) 50 mM NaCl, or (C) 150 mM NaCl. The mass for each dsRNA standard was approximated by multiplying the cited length by the average mass of a base pair (660 Da / base pair). All experiments were collected on MG-NA.

Supplementary Table S1. Fits for the calibration curves presented in Supplementary Figure S1

| Added Salt (mM) | Ionic Strength (mM) | MFP1 (protein) | | dsRNA ladder | |
|---|---|---|---|---|---|
| | | Equation | $R^2$ | Equation | $R^2$ |
| 0 | 5 | y = -0.00002709x + 0.000101 | 0.993 | y = -0.00003211 + 0.000015 | 0.999 |
| 50 | 55 | y = -0.00002369x + 0.000101 | 0.985 | y = -0.00003303 + 0.000015 | 0.999 |
| 150 | 160 | y = -0.00002491x + 0.000101 | 0.999 | y = -0.00003194 - 0.000095 | 0.999 |

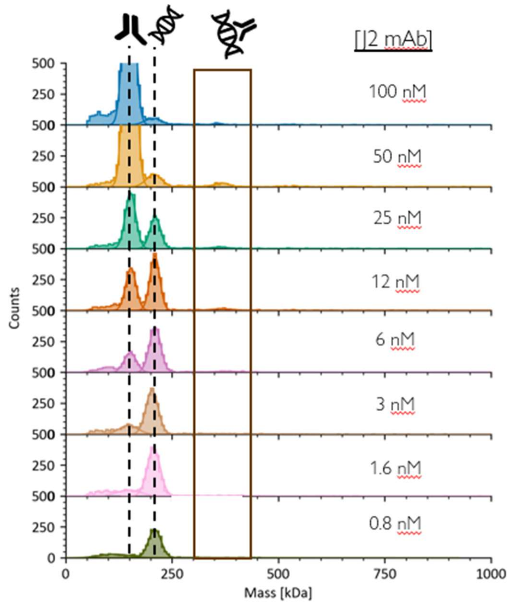


**Supplementary Figure S2. Titration o the J2 mAb at 150 mM NaCl**. Mass photometry histograms of 2 nM dsRNA substrate with increasing amounts of the J2 mAb. The free J2 mAb is observed at 150 kDa, the free dsRNA is observed at 220 kDa, and mAb/dsRNA complexes are observed above 350 kDa. Data were prepared and collected in TE buffer supplemented w 150 mM NaCl.

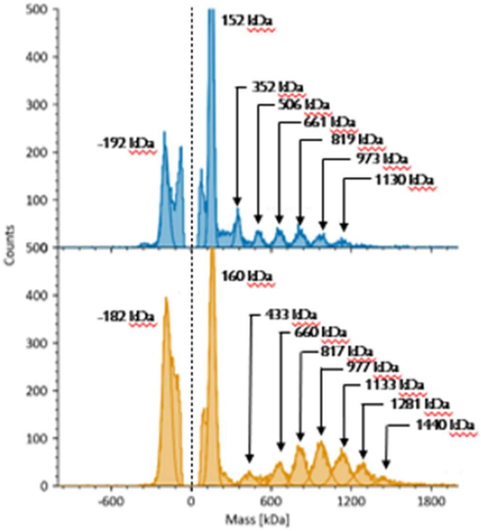


Supplementary Figure S3. Mass photometry histograms from static (top) or MassFluidix (bottom) experiments of the mAb/dsRNA complex. Samples were prepared using the J2 mAb and 400 bp dsRNA substrate at a 2:1 ratio in TE buffer with 50 mM NaCl.

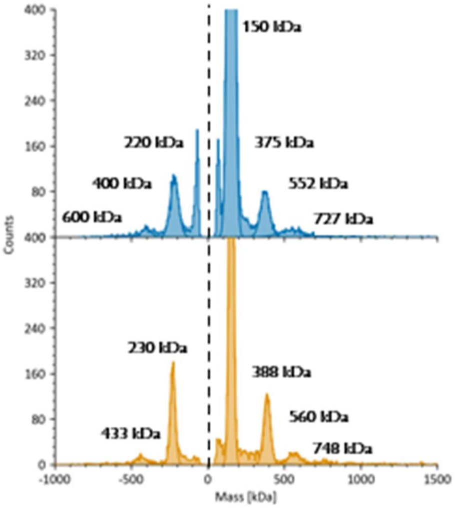


Supplementary Figure S4. Mass photometry histograms from static (top) or MassFluidix (bottom) experiments of the mAb/dsRNA complex. Samples were prepared using the J2 mAb and GFP mRNA substrate at a 2:1 ratio in TE buffer with 50 mM NaCl.
